# Demixing model: A normative explanation for inter-item biases in memory and perception

**DOI:** 10.1101/2023.03.26.534226

**Authors:** Andrey Chetverikov

## Abstract

Many studies in perception and in the working memory literature demonstrate that human observers systematically deviate from the truth when estimating the features of one item in the presence of another. Such inter-item or contextual biases are well established but lack a coherent explanation at the computational level. Here, I propose a novel normative model showing that such biases exist for any observer striving for optimality when trying to infer the features of multiple similar objects from a mixture of sensory observations. The ‘demixing’ model predicts that bias strength and direction would vary as a function of the amount of sensory noise and the similarity between items. Crucially, these biases exist not because of the prior knowledge in any form, but simply because the biased solutions to this inference problem are more probable than unbiased ones, counter to the common intuition. The model makes novel predictions about the effect of discriminability along the dimension used to select the item to report (e.g., spatial location) and the relative amount of sensory noise. Although the model is consistent with previously reported data from human observers, more carefully controlled studies are needed for a stringent test of its predictions. The strongest point of the ‘demixing’ model, however, is that it shows that interitem biases are inevitable when observers lack perfect knowledge of which stimuli caused which sensory observations, which is, arguably, always the case.

## Introduction

It is always difficult to deal with similar things. For example, it is easy to choose a place for dinner when there is a good option and a bad option, but choosing between two good ones is tricky. This is the case not only for such high-level decisions but also for perception and memory. When humans try to perceive or memorize visual features of an object and there are other similar objects present now or in the recent past (e.g., zebras in Figure 1A), their responses are not just less accurate, they are biased. Responses related to one item are shifted towards (attractive bias) or away (repulsive bias) from the others. Such inter-item biases (also known as contextual biases) permeate both perception and memory and are well known in many feature domains: size (Ben-Shalom and Ganel 2012; Mruczek et al. 2017), colour (Barbosa and Compte 2020; Chunharas et al. 2022; Kingdom 2017; Nemes et al. 2012; Lively et al. 2021; Golomb 2015; Johnson et al. 2022), orientation (Clifford 2014; J. Fischer and Whitney 2014; Rademaker et al. 2015; Yu, Rahim, and Geng 2022), spatial frequency (Huang and Sekuler 2010), motion direction (Alais, Leung, and Van der Burg 2017; Czoschke et al. 2019; Rauber and Treue 1999; Czoschke et al. 2023, 2020), etc. (see reviews in Bays et al. 2022; Chunharas et al. 2022; Pascucci et al. 2023). Even higher levels of visual hierarchy, for example, facial processing (e.g., Mallett, Mummaneni, and Lewis-Peacock 2020; Storrs and Arnold 2015; Van der Burg, Rhodes, and Alais 2019; Webster and MacLeod 2011; Kim and Alais 2021; Liberman, Fischer, and Whitney 2014; Manassi and Whitney 2022), are not free from them. But why do humans have these consistent errors? And why in some cases the differences between objects are exaggerated and underestimated in others (Figure 1C, D)? Despite a long history of research, the existing models of these biases do not provide a satisfactory computational explanation, deferring instead to physiological (“observers are biased because of the way the neurons work”) or environmental (“observers are biased because of the environment structure”) factors. Here, I argue that such biases occur because the brain is trying to solve a complicated mixture problem (Figure 1B) and are inevitable for any observer striving for optimal (the most precise) decisions, regardless of a specific neural implementation or environment.

**Figure 1:**
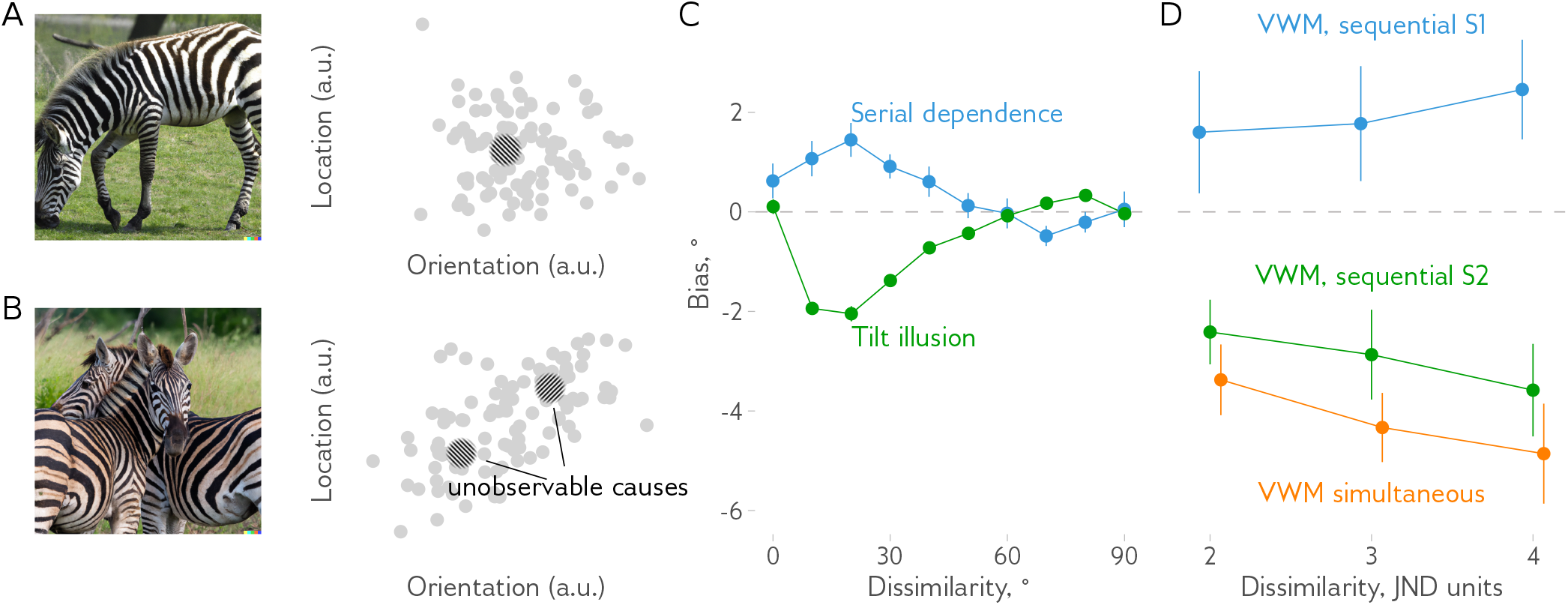
The mixture problem in perception. **A**: Consider an object, such as the zebra shown here, that can be described for simplicity as a point in a two-dimensional location-orientation feature space (shown as a Gabor patch). Due to noise in neural processing and in the environment, sensory observations (gray dots) obtained by the visual system will be scattered around the true object values. However, if enough observations are obtained, the object features can be estimated accurately by simple averaging. **B**: When there are two similar objects, such as two zebras, it is no longer enough to simply average the sensory observations, since the visual system has to also infer which object each observation belongs to. This commonly overlooked step is crucial for understanding why perceptual and memory estimates of an object’s features are often biased in presence of other items. **C**: Two such well-known inter-item biases in orientation perception are an attractive serial dependence bias (blue, data from Fritsche, Spaak, and de Lange 2020) and a repulsive tilt illusion (green, data from Moors and Wagemans 2020). **D**: In visual working memory studies, opposing biases are also well established. Reports of one of the simultaneously presented items in Czoschke et al. (2020) demonstrate repulsion (orange line), similar to a second item in the pair of sequentially shown stimuli (green line). In contrast, an attractive bias occurs when observers are required to report the first of the two sequentially shown items (blue line).

One of the well-studied domains exemplifying inter-item biases is orientation estimation. Orientation perception is biased when two stimuli with different orientations are present simultaneously or sequentially (the tilt illusion or the tilt aftereffect, (Gibson 1937; Gibson and Radner 1937; see reviews in Schwartz, Hsu, and Dayan 2007; Clifford 2014). These biases usually follow a repulsive-then-attractive pattern with similar orientations resulting in perceived orientation being shifted away from the other item, and dissimilar orientations resulting in a smaller shift toward it (Figure 1C). On the other hand, when observers have to estimate orientations of the sequentially presented stimuli, an attractive-then-repulsive pattern known as “serial dependence”^1^ occurs (Cicchini, Mikellidou, and Burr 2018; J. Fischer and Whitney 2014; Fritsche and de Lange 2019b; Gallagher and Benton 2022; Kiyonaga et al. 2017; Pascucci et al. 2019; Rafiei et al. 2023). The same pattern is observed for estimates of target orientation as a function of their similarity with distractors in a visual search task (Rafiei, Hansmann-Roth, et al. 2021; Rafiei, Chetverikov, et al. 2021). Several theories (discussed in detail later) were put forward to explain why both attractive and repulsive biases can arise but there is currently no definitive conclusion (Bliss, Sun, and D’Esposito 2017; Ceylan, Herzog, and Pascucci 2021; Cicchini, Benedetto, and Burr 2021; Collins 2020; Fritsche, Spaak, and de Lange 2020; Fritsche, Mostert, and de Lange 2017; Murai and Whitney 2021; Pascucci et al. 2019).

Just like in perception, in visual working memory studies, both repulsive and attractive biases were reported for oriented gratings (Dubé and Sekuler 2015; Ding et al. 2017; Dubé et al. 2014; Hajonides et al. 2023; Lorenc et al. 2018; Rademaker et al. 2015; Scocchia, Cicchini, and Triesch 2013; Guillory, Gliga, and Kaldy 2018). For example, Hajonides et al. (2023) asked observers to memorise two Gabor patches presented in sequence. When observers were asked to report the second item presented, their reports were significantly biased away from the first item (but not the other way around). At the same time, the reported orientations for both the first and the second items were shifted towards the item reported on a previous trial [see also Czoschke et al. (2019) for similar results in the motion direction domain, reproduced in Figure 1D]. Other studies reported attractive biases in orientation estimates even within a single trial. For example, Rademaker et al. (2015) and Lorenc et al. (2018) both presented a target to memorise followed by a distractor and found that target orientation estimates were biased towards the distractor. Visual working memory, similar to perception, shows both attractive and repulsive biases.

### Explaining inter-item biases

How to explain inter-item biases in orientation and other domains? Most of the existing ideas, especially for repulsive biases in perception, are focused on explaining the biases in terms of the properties of the neural populations involved in the processing of a given feature domain (i.e., at the mechanistic or implementation level in Marr’s (1982) classification). For example, a repulsive-then-attractive pattern for interitem effects in orientation perception has been explained with changes in tuning curves due to the spatial and temporal context in which the target stimulus is presented (see Schwartz, Hsu, and Dayan 2007 for a review). These models help to elucidate the neural basis of inter-item models and the relationship between neurophysiological changes and behaviour, but do not help to explain it at the computational level. In other words, they do not say anything about the function of the biases: why the tuning curves change in response to contextual influences or, similarly, what the utility of the observed behaviour is.

The models that addressed the question of functional relevance can be grouped into two categories. The first type invokes the idea of combining information from different sources to improve the accuracy of the estimates and is helpful for explaining attractive effects (van Bergen and Jehee 2019; Brady and Alvarez 2011; Cicchini, Mikellidou, and Burr 2018; Dubé and Sekuler 2015; J. Fischer and Whitney 2014). Such integration corresponds to the behaviour of an ideal observer (van Bergen and Jehee 2019; Cicchini, Mikellidou, and Burr 2018) who knows that stimuli in the environment are spatially and temporally correlated. For example, a cat on a table is unlikely to turn into a person except in certain wizardry schools in Scotland. Hence, it is beneficial to combine the inputs from different moments in time (e.g., the cat now and a second ago) or from neighbouring locations (e.g., patches of the cat’s fur) to determine the cat’s colours. However, when dealing with stimuli in a typical laboratory experiment that are not correlated between trials and locations, the same combination approach leads to attractive biases.

Models of the second type aim to explain repulsive biases by focusing on other environmental regularities. For example, Schwartz, Sejnowski, and Dayan (2005) described a Bayesian model for orientation estimation in the presence of a “smoothness” prior that explained to some extent biases in the tilt illusion. Notably, repulsive biases in this model arise as the smoothness prior gives a higher probability to configurations with connected edges of target and distractor (e.g., when two lines are shown, a \\ pattern will be less connected and less likely than /\ hence the perceived configuration of \| pattern is closer to the latter). For obvious reasons, this cannot be generalised to other domains (e.g., colour), or even different target-distractor configurations within the orientation domain (e.g., a target Gabor fully surrounded by a distractor Gabor patch as in Moors and Wagemans 2020). In contrast, efficient coding models arguing that the visual system aims to reduce redundancy in neural codes are highly generalisable (Clifford, Wenderoth, and Spehar 2000; Stocker and Simoncelli 2005). Such models are especially well suited for explaining the inter-item biases for sequentially presented stimuli (i.e., adaptation). In brief, they assume that the noise in the visual system is reduced for a range of features matching the first presented item (adaptor) at the expense of increased noise outside this range. This leads to an asymmetry in the likelihoods of different stimuli, which in turn creates repulsive biases in perception of the items within the vicinity of the adaptor. In combination with the Bayesian integration model described above, the Bayesian observer approach could explain both attractive and repulsive biases (Geert, Ivancir, and Wagemans (2022); see Wei and Stocker (2015) for the efficient Bayesian observer model in the context of cardinal biases, and Sheehan and Serences (2022) for a related model applied to biases in representations decoded from neural data). Note, however, that efficient coding models cannot explain repulsive biases for simultaneously presented stimuli, especially when the features of these stimuli vary randomly on a trial-by-trial basis, because it is not clear how and why this could change the corresponding neural coding mechanisms^2^. Yet, such biases are as pervasive as biases in sequential presentation, highlighting the need for a more parsimonious explanation.

### Demixing model

Here, I present an alternative approach based on the idea that inter-item biases arise because the brain does not know exactly the association of the neural responses and their causes in the external world and has to infer it. In general, the “inverse inference” problem is well known in vision science (e.g., Kersten, Mamassian, and Yuille 2004; Pizlo 2001; Rock 1997). Different items can create identical sensations (for example, different 3D objects can look identical when projected on the 2D plane), while the same item can look differently under changes in illumination, distance, etc. The brain then needs to infer what objects are based on noisy sensory observations (Figure 1A). What is often overlooked is that to infer the object properties, such as the location or orientation of a Gabor patch, the visual system has to infer which objects caused which observations as well. This is especially problematic when similar objects are presented close in time or space, that is, under the conditions in which the inter-item biases are usually observed. The key proposition of the current work is that this problem can be seen as a mixture problem: the observer is presented with a mixture of sensory observations and has to determine the properties of the distributions that generated them (or ‘demix’ the observations). In the following, I show that an optimal solution for this problem is biased when the items are similar and sensory noise is high. Furthermore, the nature of the biases (e.g., attraction or repulsion) depends on the relative amounts of sensory noise and the assumptions the observer has about the structure of the environment.

To explain the model, I will start with a toy example showing how demixing the observations can lead to biases in decisions when the stimuli are described on just one feature dimension (e.g., orientation or colour). I will then show the optimal solution for this problem and will prove that it is also biased. Then, I will consider a more common case of a two-dimensional mixture problem (e.g., orientation combined with location). I will then describe the model predictions for a very interesting case of unequal stimuli noise and show that in such cases opposing repulsive and attractive biases are expected within a given trial.

#### A toy example

To illustrate the intuition behind the model, consider first a toy example with two stimuli generating two sensory observations each in a single feature dimension, such as colour or orientation. The sensory observations are noisy, so each stimulus can lead to different observations. The probability with which each stimulus generates a given observation can be described as a probability distribution (a *generative* distribution). If the stimuli are similar, for example, one is only slightly redder than another or has only slightly different orientation, their respective distributions will overlap (Figure 2A), so many observations could be caused by either stimulus. In contrast, when they are different, there will be little overlap between their corresponding distributions (Figure 2B).

**Figure 2:**
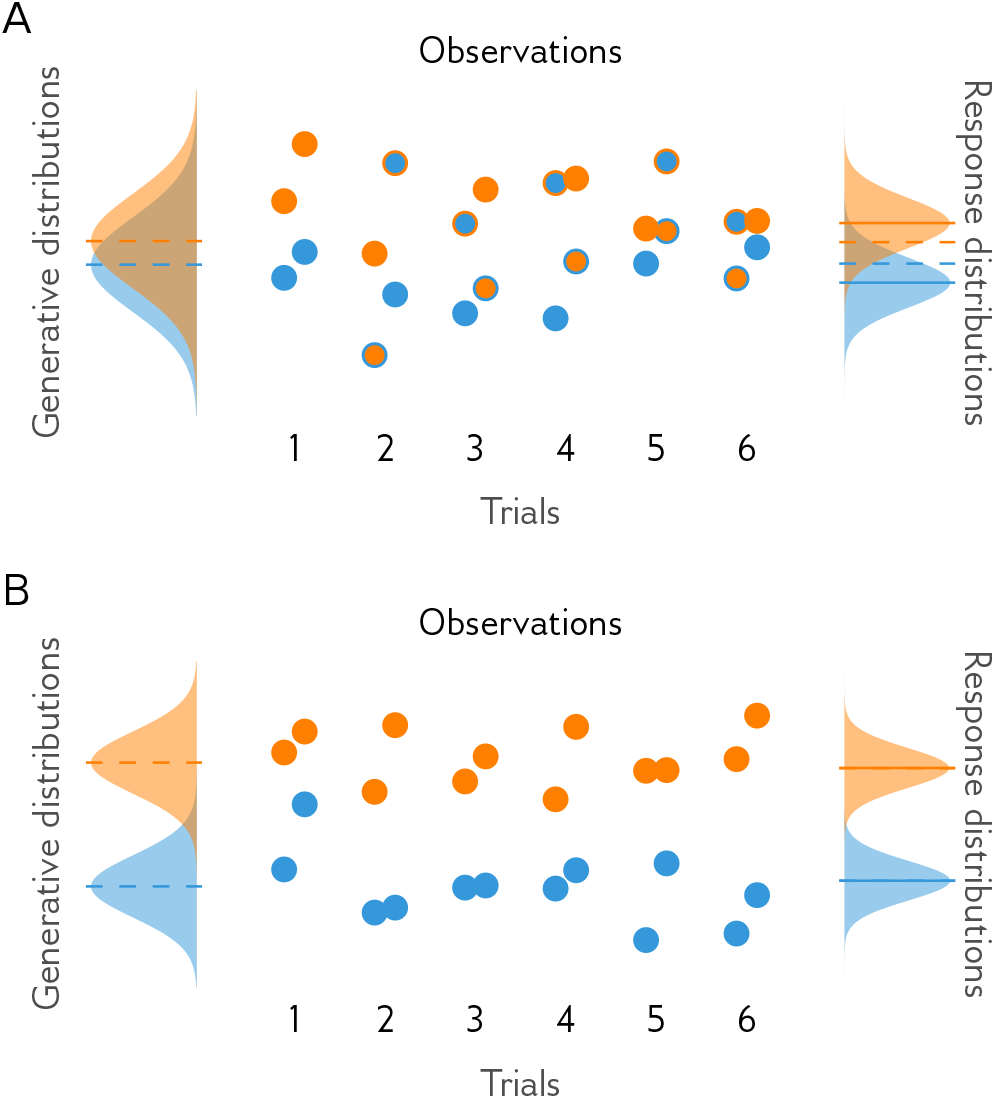
An illustrative toy example. **A**: Consider two stimuli with similar feature values (e.g., colours or orientations) indicated by dashed lines in the left panel. Due to sensory noise, the values of sensory observations sampled from these two stimuli can be represented as two overlapping probability distributions (generative distributions, left panel). An observer samples two observations from each of the stimuli in several trials (middle panel) and assigns each of the observations to one of the stimuli. The actual stimulus from which the observation was sampled (inner colour of dots) and the assigned one (their outer colour) often mismatch because similar observations are likely to be grouped together. The stimuli feature values are then estimated by averaging the observations assigned to each stimulus. Bottom: As a result, the response distributions (right panel) differ from generative distributions and the means of the distribution of estimated stimuli features on average (solid lines) the responses will be shifted away from the true feature values (dashed lines). **B**: Same as in A but with more dissimilar stimuli. In this case, the observations are likely to be assigned correctly to the stimuli that caused them, and the average of the response distribution matches the stimuli.

The observer wants to infer the stimuli properties, that is, the feature values corresponding to the mean of the distributions, but is unaware of the true generative distributions and have to determine their parameters based on the observations obtained. I assume for the simplicity of this toy example that the observer correctly expects that each stimulus generates the same number of observations (the sensory sampling rate is the same for the two stimuli).

It is intuitively clear that if the four observations are to be split in two pairs attributed to two unknown stimuli, the pairs would be formed using the samples that are close by. This is because the observations that are very similar are more likely to be caused by the same stimulus different from each are less likely than the ones that are similar different (assuming for any unimodal noise distribution). Then, for example, the pair with the redder observations is attributed to stimulus A, and the pair with the yellower observations to stimulus B. When the same stimuli are encountered the next time, the pairs are again formed from the closest observations, and one of the stimuli is yellower than the other. So each time, the difference between stimuli is exaggerated, because the redder samples from the yellower stimuli are likely to be attributed to the redder stimuli and vice versa. This toy example illustrates the intuition behind the repulsive bias in the ‘demixing’ model: When stimuli are similar, the sensory samples from one stimulus are often attributed to another, leading to a bias away from that stimulus.

However, this toy model is suboptimal because it does not account for uncertainty of the assignment of a given observation to a stimulus. That is, when the observations are similar to each other, it is difficult to say for sure which stimulus caused which observation and that uncertainty is important for the observer to make accurate predictions. A more optimal model is then needed.

#### The optimal solution for the one-dimensional mixture problem

The optimal solution to the mixture problem is to estimate the probability that each stimulus caused a given observation instead of assigning them with 100% certainty as in the toy example above. This is a well-known problem in statistics, but the solution to it is complex. Consider a more general case where a set of *N* observations **x** is drawn with a probability of *λ* from one of the two normal distributions with means *μ*_1_ and *μ*_2_ and standard deviations *σ*_1_ and *σ*_2_. The probability of observing **x** is determined as:

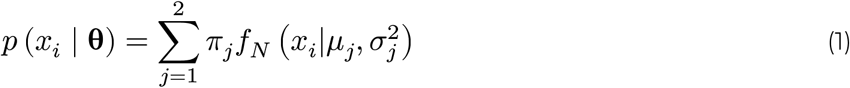

where **θ** = {*μ*_1_, *μ*_2_, *σ*_1_, *σ*_2_} is a set of parameters describing this mixture distribution, *π*_*j*_ are their weights in the mixture **π** = {*λ*, 1 − *λ*}, *j* is the component index, *i* is the observation index, and *f*_*N*_ is the normal distribution probability density function. The likelihood of a certain **θ** is then given by:

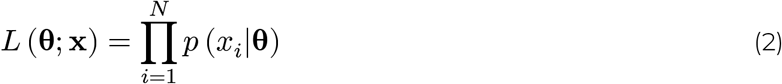

where *N* is the number of observations.

Finally, the posterior distribution can then be found following the Bayes rule:

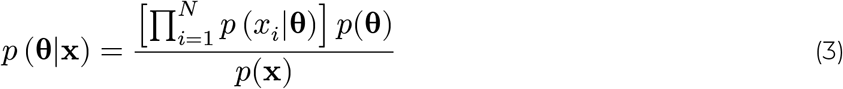

Assuming that there is no prior information about distribution parameters, their best estimates can then be found by maximizing the likelihood.

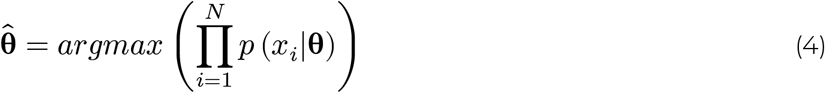

This is the optimal solution for the one-dimensional mixture problem as it maximizes the probability of getting the correct estimate. Notably, this likelihood gives to 2^*N*^ terms when expanded, making exact computations time-consuming. Luckily, there are well-known algorithms, such as the expectation-maximization (EM) method used in the simulations described below, that allows finding the solution for a given sample of observations quickly.

#### The simulations for a one-dimensional mixture problem reveal a repulsive-then-attractive bias pattern

I simulated the optimal (maximum likelihood) estimates of the means for the two distribution case assuming that the stimuli are equally noisy (*σ*_1_ = *σ*_2_) and generate equal number of observations (*N* = 100). The simulations were done for 10000 trials with different amounts of noise (*σ* ∈ {5, 10, 20, 30}, arbitrary units, a.u.) and different degrees of similarity between the stimuli (|*μ*_1_ − *μ*_2_| ∈ [0..120]). For each trial, the most likely solution was determined by the EM algorithm using *Rmixmod* package in *R* (Lebret et al. 2015). The error of the estimates was then computed as a difference between the true and the estimated means. The bias corresponds to the error with the error sign flipped in such a way that a positive sign indicates that the estimated mean is shifted from the true mean for the corresponding mixture component towards the other component mean (an attractive bias) while the negative one corresponds to a shift away from the other mean (a repulsive bias).

As shown in Figure 3A, I observed the repulsive biases for similar stimuli, however, I also found a weaker attractive bias for dissimilar stimuli. The attractive bias switches to repulsive when the similarity (the difference between the mean feature values of the stimuli *μ*_1_ and *μ*_2_) is about 0.8 of the noise amount. Such repulsive-then-attractive pattern is common for inter-item biases in perception literature, such as the tilt illusion exemplified in Figure 1C.

**Figure 3:**
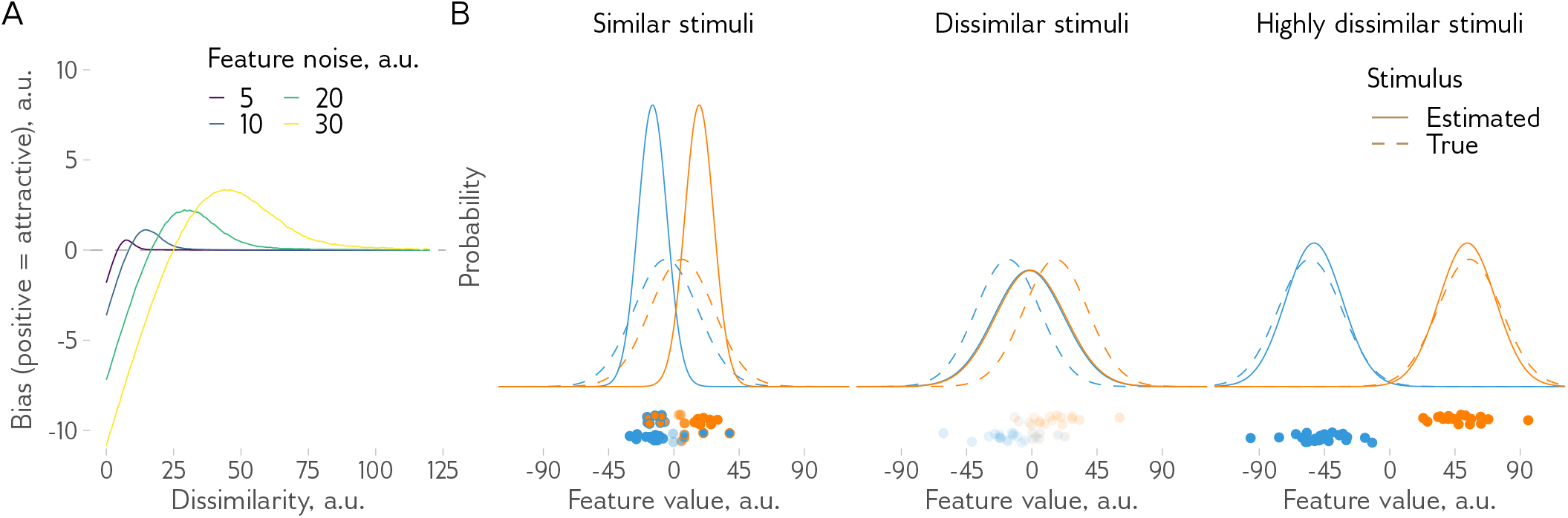
A simplified one-dimensional mixture problem (e.g., a stimulus is defined by orientation only). **A**: The average bias for a one-dimensional mixture problem as a function of stimuli dissimilarity and the amount of noise. For all noise levels, the model shows a repulsive-then-attractive pattern. **B**: Example cases for different dissimilarity levels. Dashed lines show the generative distributions, and solid lines show the distributions based on the estimated parameters, 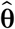. Dots show sensory observations sampled from the generative distributions (dot’s inner colour indicates the generative distribution it was sampled from, the outer colour - is the assigned distribution, transparency corresponds to the confidence in the assignment, that is, the deviation of weight in the mixture from 0.5). With highly similar stimuli, the mixture problem is resolved by distributing the observations into well-separated groups, which leads to a repulsive bias.

Similarly to the toy example discussed above, the repulsive bias arises because observations in a mixture are grouped together by similarity. Only in this case, the grouping is probabilistic (i.e., a given observation has a certain probability of belonging to each of the groups; see Figure 3B, left panel). This is, in turn, because similar observations are more likely to come from the same source compared to dissimilar ones^3^. Hence, observations that are far from the true stimulus and close to another stimulus are likely to be associated with the latter. So both stimuli ‘lose’ the observations that are close to the other stimuli, so they look less similar to each other. That is, the dissimilarity between stimuli is overestimated, the repulsive bias occurs.

Why does the attractive bias arise? The analysis of the simulated data revealed that they stem from trials in which the probabilities of belonging to each group assigned to the observations were close to 0.5, as exemplified by Figure 3B (middle panel). This represents another solution to a mixture problem. Instead of assigning the observations to one of the sources with high probability, the model assigns probabilities closer to 0.5, essentially suggesting that stimuli are the same.

Of course, when stimuli are very dissimilar, there is no ambiguity as to which stimulus caused which observation, and the responses are unbiased Figure 3B (right panel). However, as long as ambiguity exists, both solutions are possible and the average bias (Figure 3A) depends on their relative balance (Figure S2).

#### Two-dimensional mixture problem

The problem described above does not yet make justice to the problem that observers typically have in experiments described in the introduction. The reason is that in a typical experiment, there are two dimensions present, namely, a response dimension and an identifying dimension. The response dimension is the dimension on which the response is made while identifying dimension is the feature dimension determining to which stimuli the observer has to respond. For example, in the case of a tilt illusion, an observer might need to estimate the orientation (the response dimension) of a grating presented at the centre of the screen while a surrounding grating needs to be ignored (the location being an identifying dimension). Similarly, in a typical serial dependence study, observers need to estimate the orientation of currently presented stimuli while the previous one has to be ignored (the time being an identifying dimension). In general, participants have to report a certain feature while space, time, or another feature serves to identify the relevant item.

When treated formally, this task can be described as a two-dimensional mixture problem. That is, instead of univariate feature distributions from the previous sections, sensory observations can now be described as bivariate distributions (Figure 4A). The goal of the brain, however, remains the same: to infer the properties of the underlying distributions without knowing which observation is caused by which stimulus.

**Figure 4:**
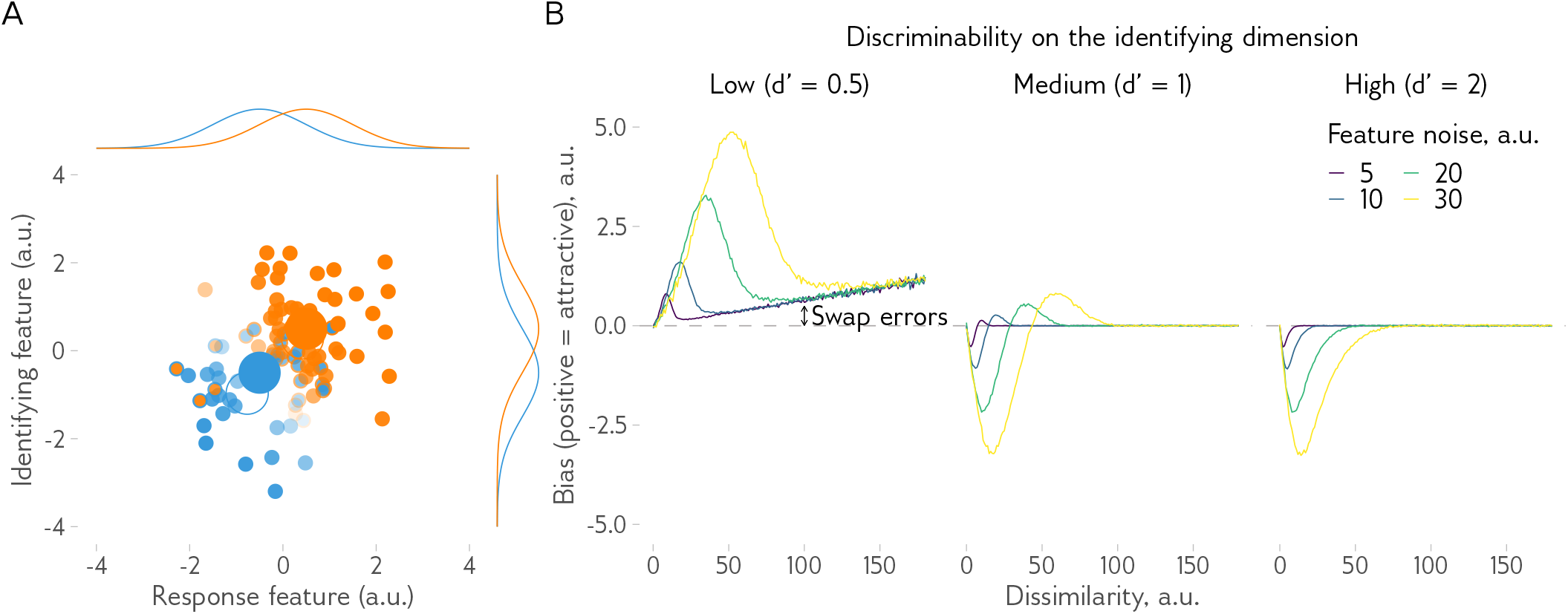
A two-dimensional mixture problem (e.g., a stimulus is defined by a combination of orientation and spatial location). **A**: An example of a two-dimensional mixture problem and a probabilistic solution. Two stimuli (solid circles) are used to generate sensory observations (dots) with noise on the two dimensions described by an independent bivariate probability distribution shown with side plots on the top and on the left. A demixing model is then used to estimate the most likely stimuli properties (outlined circles). The dot fill colour shows the distribution from which they were generated with the colours matching the stimuli, and the dot outline colour shows the stimuli to which they are more likely to be assigned by the model with less transparent dots indicating higher certainty in the assignment. **B**: The average bias in response distributions for a two-dimensional mixture problem. Each line shows an average bias as a function of dissimilarity between stimuli (on the response dimension), while discriminability (on the identifying dimension) varies between panels. colours indicate the amount of feature noise on the response dimension.

What is the optimal solution for a two-dimensional mixture problem? I again used simulations to answer this question. The feature similarity was varied as before, and for the identifying dimension, I consider three cases differing in discriminability defined as a *d*^′^ measure:

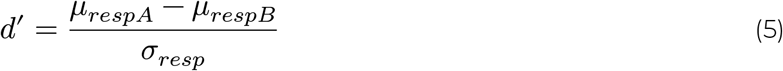

where *μ*_*respA*_ and *μ*_*respB*_ are the means of the two stimuli distributions in the response dimension and *σ*_*resp*_ is their standard deviation (the same for the two stimuli). Three levels of discriminability were used: low (*d*^′^ = 0.5), medium (*d*^′^ = 1), and high (*d*^′^ = 2). For each discriminability level, I then simulated the optimal solutions in 10000 trials for different levels of feature noise and dissimilarity.

Notably, for the medium dissimilarity level, the biases showed the same repulsive-then-attractive pattern as in the 1D case (Figure 4B). In contrast, low discriminability on the identifying dimension led to an attractive-only pattern, while in the high-discriminability condition, the repulsive-only pattern was evident. The repulsive pattern arises due to the same reasons as in the one-dimensional case. Namely, the observations are more likely to be associated with causes (i.e., stimuli) that are closer to them. Hence, the observations between the two stimuli in sensory space are not always associated with the stimulus that caused them, leading to a bias. But why does the strong attractive pattern appear in the low discriminability condition?

The attractive pattern in the low-discriminability condition comes from a combination of two effects. First, the model often converges on a solution with the distributions centred around the average of the two stimuli (similar to Figure 3B, middle panel). Secondly, the associations between the response and the feature dimensions are often incorrect, leading to “swap errors”. The latter is also responsible for a pattern of increasing bias at large feature distances in the low-discriminability condition, because the same share of swap errors corresponds to a larger bias in the average feature estimate when the distributions are further away from each other.

Notably, in high-discriminability conditions, only repulsive bias is observed on average, because a solution with very similar stimuli is no longer likely as they clearly differ on identifying dimension (e.g., location). In sum, the simulations for the two-dimensional mixture problem show that different bias patterns are possible, from attractive-only to repulsive-only, providing a potential explanation for a diverse pattern of results observed in experiments with human observers.

#### Unequal noise: the effect of memory decay, attention, and other factors

So far I assumed that the stimuli have equal noise levels. This is, of course, not always so in reality. For example, when two stimuli are presented consecutively, the second stimulus appears while the first one is held in memory, so arguably the first one would have higher uncertainty. So unequal noise levels are to be expected in the typical conditions of serial dependence studies or a memory task with sequentially presented items. Similarly, if the items are presented simultaneously, they also would not have equal noise levels because of the many factors affecting the distribution of attention. For example, if an item is similar to a previous target or located at the same place, it is likely to be attended more than the other item, hence it would have lower noise levels. What are the model predictions in such cases?

I simulated the optimal solutions for the unequal noise and medium discriminability case 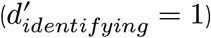. A higher noise level varied from 5 to 60 a.u. (*σ*_*higher*_ = {5, 10, 20, 30, 60}) while the lower noise level varied from 5 to 30 a.u. (*σ*_*lower*_ = {5, 10, 20, 30} with *σ*_*higher*_ > *σ*_*lower*_). Figure Figure 5A shows the bias for the two items averaged over the higher-noise item levels as a function of the lower-noise item noise. Interestingly, the lower-noise item always shows an attractive bias while the high-noise item shows a repulsive bias (this pattern holds in general for other discriminability levels but there are some small exceptions, see Figure S4 for details).

**Figure 5:**
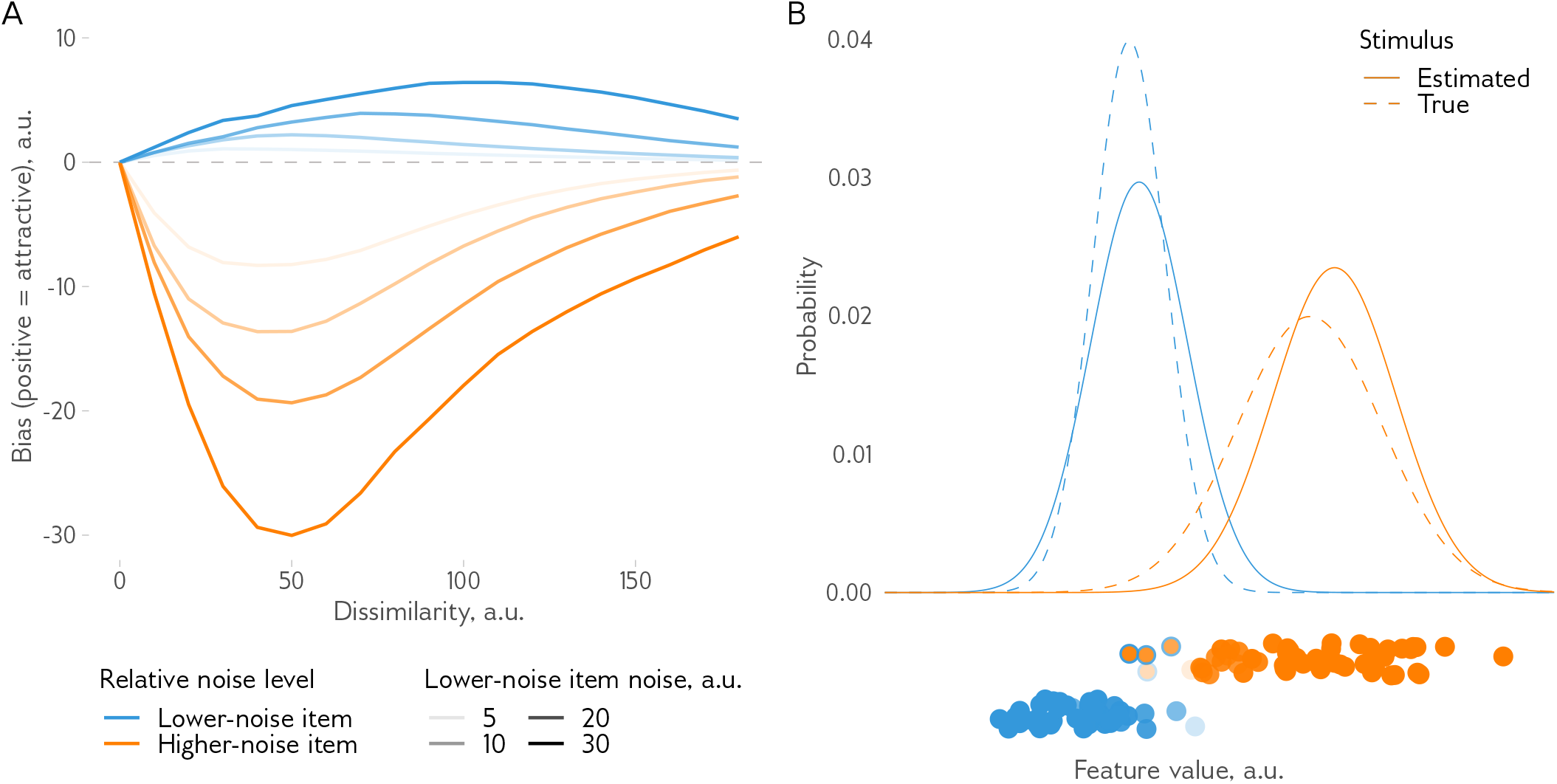
Demixing model predicts a diverging pattern of biases for items with unequal noise levels. **A**: On average, the lower-noise item (blue) is attracted to a high-noise item, while the higher-noise item (orange) is repulsed from the lower-noise one. The strength of the bias depends on the exact amounts of noise (shown here for the lower-noise item in different lines), but the diverging bias pattern stays the same. **B**: An example solution for the unequal noise two-dimensional mixture problem (only the response dimension is shown for simplicity). Each stimulus can be characterised as a probability distribution on the dimension of the response feature. Compared to the true stimuli features (dashed lines), the estimated features (solid lines) are shifted, the stimulus with the lower noise (blue) is shifted towards, while the stimulus with the higher noise (orange) is shifted away from the other stimulus. The bottom panel shows the sensory observations with their true generative distribution shown as the fill colour and the most likely assigned distribution as the outline colour. The transparency of the dots shows confidence in the assignment. The observations from the higher-noise distribution that is close to the lower-noise distribution are ‘captured’ by the latter (orange dots with the blue boundary). This shifts the lower-noise estimated distribution closer to the higher-noise one, while the latter, after ‘losing’ the observations on one side is moved in the opposite direction, away from the lower-noise distribution.

Why does it happen? Figure 5B shows an example solution for a two-dimensional mixture problem with unequal noise, focusing on the response dimension. The tail of a high-noise distribution overlaps with the low-noise distribution, thus the observations from that tail are likely to be assigned to the low-noise distribution. The low-noise distribution will then become attracted to the high-noise distribution. The high-noise distribution, on the other hand, will be repulsed away from the low-noise distribution after “losing” the observations from the tail on one side.

## Discussion

This paper introduces the demixing model to explain inter-item biases in perception and visual working memory. Both repulsive (perceived stimuli features are shifted away from the other stimulus) and attractive (toward the other stimulus) biases are well known in the literature for different features including colour, size, motion direction, or orientation (e.g., Ben-Shalom and Ganel 2012; Mruczek et al. 2017; Barbosa and Compte 2020; Chunharas et al. 2022; Kingdom 2017; Clifford 2014; J. Fischer and Whitney 2014; Rademaker et al. 2015; Alais, Leung, and Van der Burg 2017; Czoschke et al. 2019; Rauber and Treue 1999). However, there is a notable lack of theoretical models able to explain this variety at a computational level in Marr’s (1982) classification. In other words, existing models do not provide a clear answer as to why the biases exist. The models combining efficient coding and Bayesian integration principles (Sheehan and Serences 2022; Fritsche, Spaak, and de Lange 2020) come close but cannot explain the repulsive effect arising in simultaneous presentation. The ubiquity of the inter-item biases suggests that general computational principles explaining such biases should exist.

The demixing model explains both attractive and repulsive inter-item biases as an inevitable consequence of the simple fact that the brain cannot know in advance what causes the neural responses evoked by stimuli and has to infer these causes. When there is just one stimulus or they are very distinct, it is not an issue. However, when there are two or more similar stimuli involved, the indeterminacy of the link between the stimuli and the observations poses a problem. What I show here is that the optimal solutions to this problem that maximize the probability of getting the correct estimate are biased.

When there are two stimuli defined by a combination of two features, one is reported (the response dimension, e.g., orientation or colour) and another is used to determine which stimulus to report (the identifying dimension, e.g., location or time). The direction and strength of the biases depend on multiple factors, of which here I considered two: (1) the similarity between items; (2) the amount of sensory noise along the response dimension; and (3) the discriminability of the stimuli on the identifying dimension.

The results show that similarity affects the bias strength. This is not surprising and matches a well-established pattern in the literature of biases gradually increasing and then decreasing in strength with increasing dissimilarity (see, e.g., Figure 1). The model, however, makes novel predictions for the other two factors considered. Sensory uncertainty along the response dimension should primarily modulate the strength of the biases, with the estimates for more uncertain stimuli being more biased. Discriminability on the identifying dimension affects both the strength and the direction of biases. A notable caveat is that sensory uncertainty does not determine the bias direction only when stimuli have equal noise levels in the response dimension. However, when they are not equal the biases are no longer in the same direction. The more noisy item is repulsed away from the less noisy one, while the latter is attracted to the former.

Notably, the model presented here is a normative model, which means that it describes the optimal solution to a given problem. The particular method with which this solution is found will not change the results of the model. Furthermore, less optimal models, such as, for example, a clustering algorithm (e.g., the one implemented in the toy example), linear classifiers, and other models aimed at the same goal, should exhibit similar biases (see Figure S3 for examples). The basic insight of the model, the inevitability of biases, would stand regardless of its specific implementation. And even if future studies would show that some of the model predictions are incorrect, the deviations from these normative predictions will help to guide the search for the mechanisms of working memory and perception (Geurts et al. 2018).

### Predictions for empirical studies

These model predictions need to be tested on empirical data. The effect of sensory uncertainty is consistent with recent data on biases in VWM between simultaneously presented items. Scotti et al. (2021) and Chunharas et al. (2022) found that repulsive biases become stronger with an increasing delay period before the report, corresponding to increasing memory noise. Interestingly, Lively, Robinson, and Benjamin (2021) found that repulsive biases can turn into attractive ones as the working memory load increases. This seems to contradict the results described above and the model predictions. Note, however, that working memory load could affect discriminability not just along the response dimension but along the identifying dimension as well. This could lead to a switch from repulsion to attraction as shown in Figure 4. The data from studies with a sequential presentation design, particularly in the ‘serial dependence’ also shows that the attractive bias towards the previous item increases with sensory noise manipulated through factors such as attention or contrast or sensory uncertainty decoded from neural data (e.g., van Bergen and Jehee 2019; Rafiei, Chetverikov, et al. 2021; Ceylan, Herzog, and Pascucci 2021; Cicchini, Mikellidou, and Burr 2018; Gallagher and Benton 2022; J. Fischer and Whitney 2014; Fritsche and de Lange 2019b; see Pascucci et al. 2023 for review). Note, however, that the effects of noise in sequential designs should be modelled and studied more carefully to account for the fact that by the time the second stimulus appears, there is already some representation of the first item. In contrast, the results reported here were simulated under the assumption that both items generate an equal number of sensory observations and no prior information is available. The simultaneous presentation as a simpler case presents a more obvious testing ground for the predictions described here.

The effects of discriminability along the identifying dimension on biases have also not been studied in detail. In one recent study, Yu, Rahim, and Geng (2022) found that when participants are probed for their memory of a visual search target, their responses are biased away from distracting items more along one feature dimension (colour), when they are less discriminable along another dimension (orientation), in line with the model predictions. In perception, the separation between the test item and the surrounding context is known to decrease the magnitude of the tilt illusion. Qiu, Kersten, and Olman (2013) found that the tilt illusion reduces when the test item has different contrast or visual depth compared to the surrounding inducer item. However, I am not aware of any studies that showed a complete reversal of the illusion, which could be because in all cases the test item and the inducer have high spatial discriminability. In tilt aftereffect studies, on the other hand, short delays between the adapter and test stimuli with briefly presented test stimuli can result in an attractive tilt aftereffect (Quiroga, Morris, and Krekelberg 2019). Interestingly, in motion perception, the perceived separation between the directions of motion of two overlapping random dot clouds changes from overestimation (repulsive bias) to underestimation (attractive bias) as motion coherence decreases (Gaudio and Huang 2012). However, in this study, the discriminability along the identifying dimension and the noise on the response dimension are confounded, making interpretation difficult. In VWM studies, biases tend to decrease with increasing delay between the two items presented sequentially (Czoschke et al. 2020). Similarly, C. Fischer et al. (2020) showed that an attractive bias between trials (‘serial dependence’) decreases when targets differ on identifying dimension (colour or serial position). These findings are in line with the model predictions, but more stringent tests have to be done in future studies.

Finally, the model treats space and time similar to other features, such as orientation or color. So the biases should also be observed in these domains and indeed there is some evidence showing repulsion and attraction, for example, for spatial locations (e.g., Almeida, Barbosa, and Compte 2015; Van der Stigchel et al. 2007; Schutte, Keiser, and Beattie 2017; Guillory, Gliga, and Kaldy 2018; Schutte and DeGirolamo 2020). However, the most interesting prediction of the demixing model in this context is that noise in space and time and noise in other features should interact. Similarly to how spatial discriminability affects the strength of orientation biases as discussed above, orientation discriminability should affect spatial biases. This is in contrast to models that treat space or time as something special and different from other features. For example, in serial dependence studies, the idea of a spatial ‘continuity field’ introduced by J.

Fischer and Whitney (2014) suggests that the serial dependence effect should diminish as the spatial distance increases between the inducer and the test item. The demixing model predictions agree to some extent, but also suggest that the effect of spatial distance should depend on the feature noise (Figure S6) and can change from attraction to repulsion. The interdependence of different features (including space and time) is an important characteristic of the demixing model that leads to unique predictions for empirical data.

### Limitations

The model described here assumes that the observer uses a model that matches the generative model and treats the observations as a mixture of two stimuli. This is in contrast to, for example, standard Bayesian observer models of inter-item biases such as serial dependence (van Bergen and Jehee 2019; Cicchini, Mikellidou, and Burr 2018). In other words, this a demixing model, not a mixing one. But this is of course not a given. Even if a participant in an experiment is instructed to discriminate two stimuli, the visual system might not care about the instruction. In the extreme, when two stimuli are identical, why perceive them as different? In other words, the brain might infer not just the stimuli features but the number of stimuli as well. For an optimal observer, the inferred causal structure (i.e., the number of stimuli behind the sensory observations) depends on the relative amount of evidence for the two models. Notably, a model with more hypothesized stimuli would always fit data better (Figure S5), so the model complexity should be penalized. However, it is not easy to determine which penalty should be used and whether it should be a flexible or a fixed one, so it is outside of the scope of this paper.

I do not suggest that demixing is the only source of repulsive biases in the common perceptual and memory tasks. For example, when two stimuli are separated by the categorical boundary (e.g., two Gabor patches are oriented at +5 and -5 degrees from horizontal orientation), repulsive effects can be exaggerated due to anisotropies in encoding in line with efficient coding models or due to other, postperceptual processes (Harrison, Bays, and Rideaux 2023; Wei and Stocker 2015; Taylor and Bays 2018; Fritsche and de Lange 2019a). Similarly, the model does not include any priors for simplicity, while a strong prior can lead to results that could be seen as attractive bias (i.e., the difference between items on the opposite sides of a prior would be underestimated). A relative contribution of different bias sources to the magnitude and direction of biases remains to be seen. However, one could also argue that such effects are not really inter-item biases and they should be accounted for through careful study planning when the goal is to understand specifically how one item influences another.

### Conclusions

The demixing model presents an alternative take on inter-item biases in perception and working memory. Compared to classic ideal (Bayesian) observer models and efficient coding models, this model assumes that each stimulus generates multiple observations and the observer does not know the stimuli generating them but has to infer them from the data at hand. Then, through computational modelling, I show that the optimal solution for this problem is biased with different bias patterns depending on the task parameters. I leave the tests of the model outside of the scope of this paper, as the model presented here is a normative one and as such its predictions about what the observer *should* do are the same regardless of what real observers are doing. However, it is promising that such a simple model provides patterns of results surprisingly similar to previously observed data.

## Acknowledgments

I am grateful to Sabrina Hansmann-Roth and David Pascucci for comments on the early versions of the manuscript.

## Data and code availability

The data and code for the simulations reported in this paper are available at: https://osf.io/mbhsg/?view_only=1a57a57c4188499f8d510433c8038564.

## Supplementary figures

**Figure S1:**
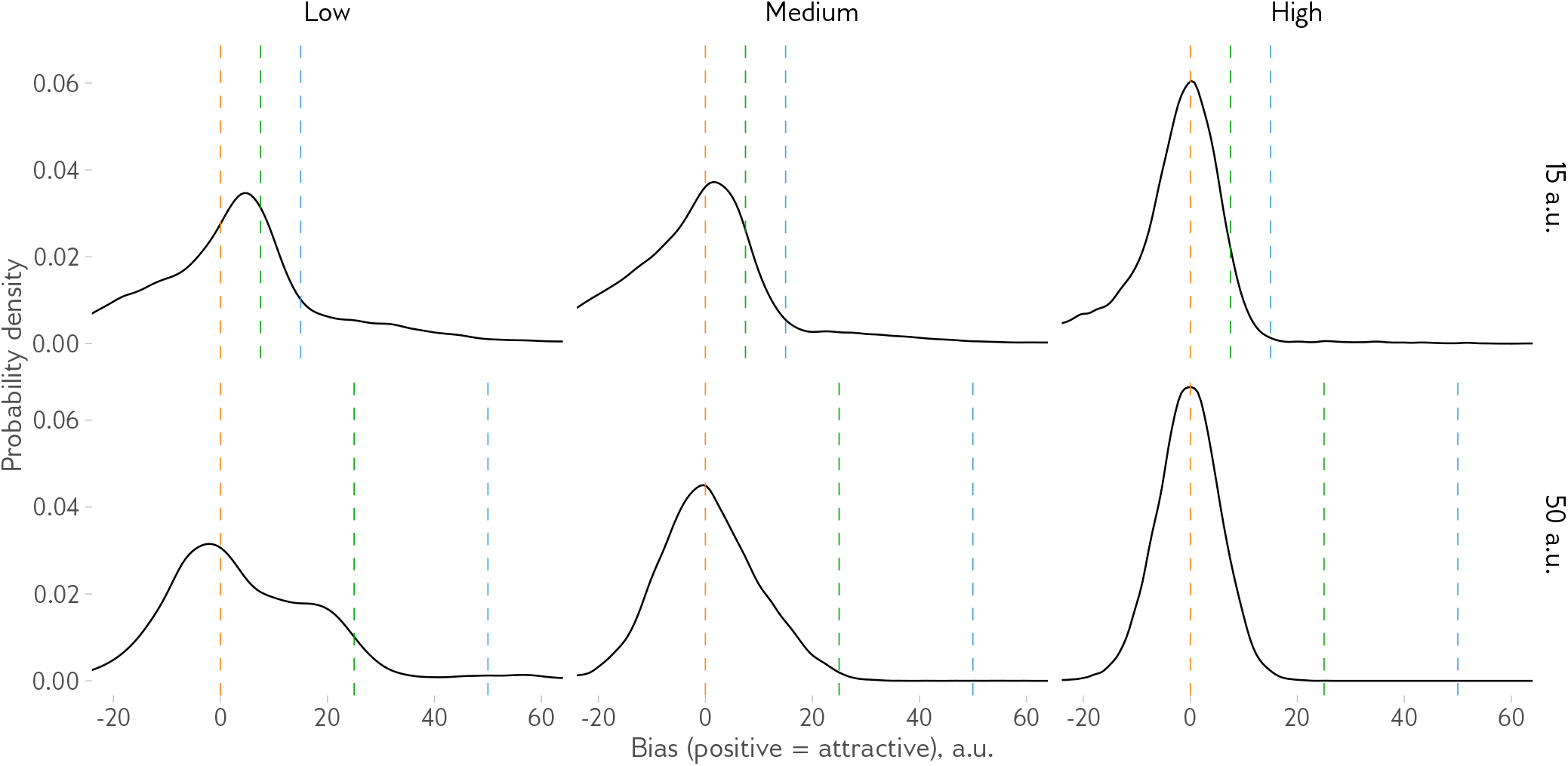
Model response probability distributions. To further understand the model responses, I analysed the response distributions for the 2D mixture model with different discriminability along the identifying dimension (low, medium, or high, in facets; the discriminability is defined the same way as for Figure 4 with *d*^′^ of 0.5, 1, or 2 on the identifying dimension, respectively) at a given dissimilarity level of 15 or 59 a.u. Dashed lines indicate the true mean of a given stimulus (orange), the midpoint between stimuli (green) and the other stimulus (blue). In the high-discriminability condition at 50 degrees a.u. dissimilarity, the response distribution is unbiased and perfectly symmetric. When the dissimilarity is low, the distributions are asymmetric, with more responses with negative (repulsive) bias. In addition, with low and average discriminability, a long tail on the right is noticeable, corresponding to swap errors. Finally, when discriminability is low and dissimilarity is high (bottom-left), many responses are close to the average of the two stimuli, matching a strong attractive bias visible in the average bias plots (Figure 4).

**Figure S2:**
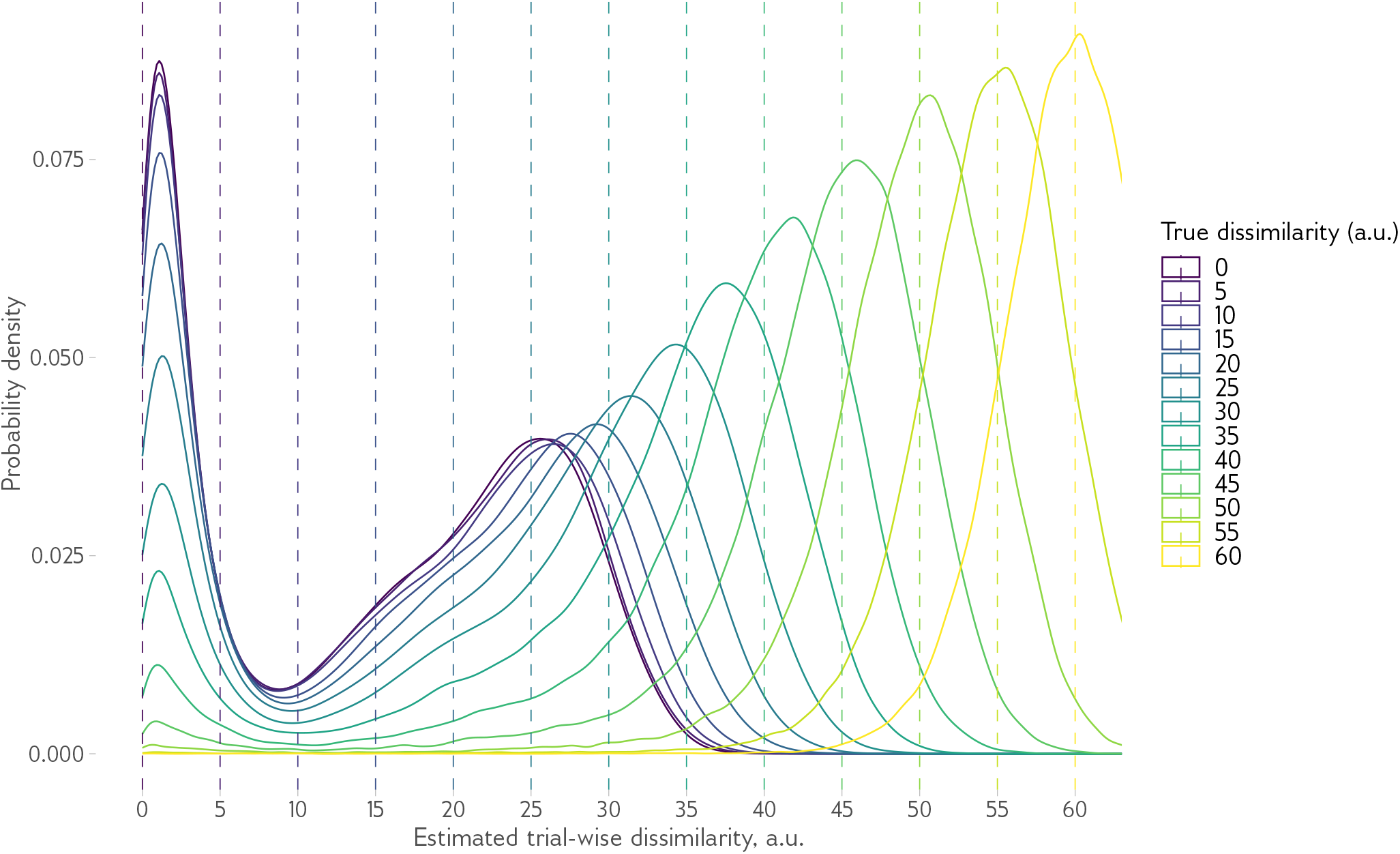
Estimated dissimilarity for the one-dimensional demixing model. Each line shows the probability of observing a certain dissimilarity in the model solutions as a function of true dissimilarity between stimuli. Both a solution with almost identical stimuli (the peak around zero) and the solution with overestimated dissimilarity is clearly visible for all levels of true dissimilarity up until about 40 a.u. However, the amount by which the dissimilarity is overestimated (that is, the difference between the true and the estimated dissimilarity) is reduced with increasing true dissimilarity. Accordingly, the average bias (Figure 3) changes from repulsion to attraction to no bias as dissimilarity increases. Dashed lines show the true dissimilarity level. The feature noise is 20 a.u. for both stimuli in this figure.

**Figure S3:**
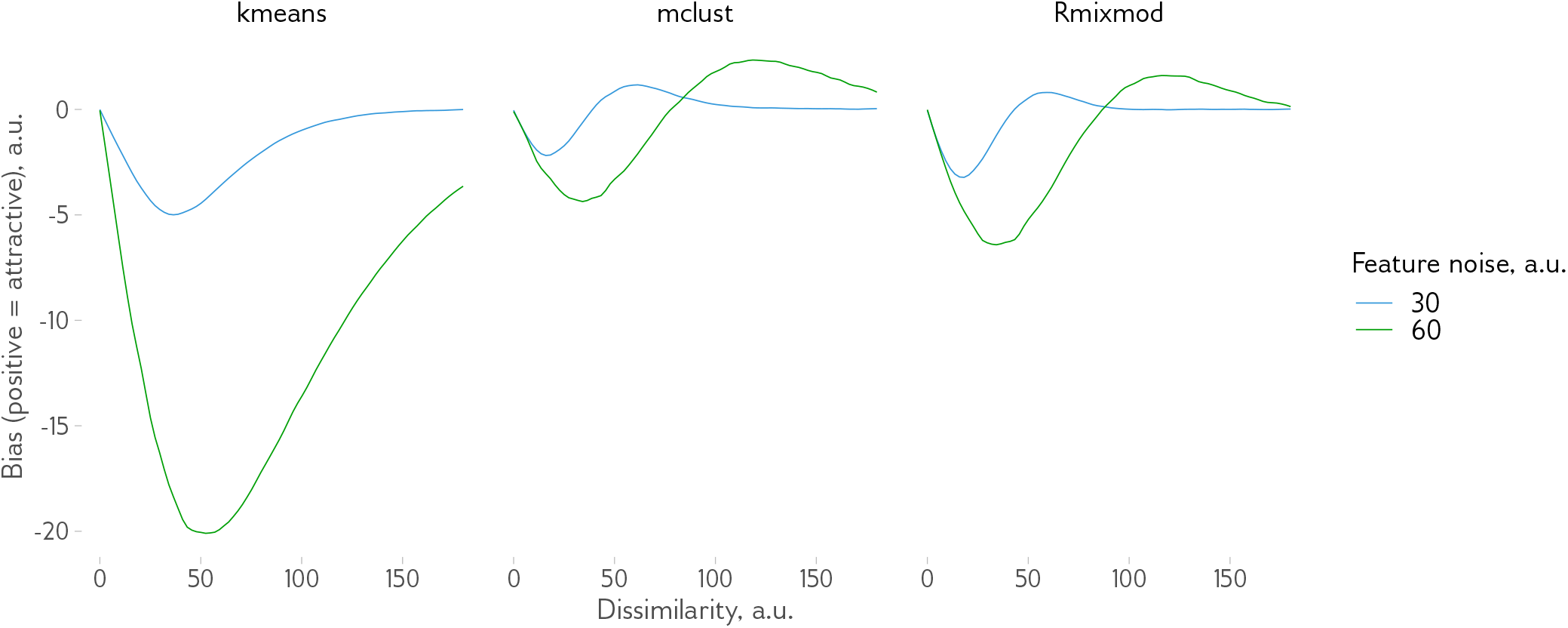
Comparing different methods. To test the robustness of the results, I compared two different implementations of the expectation maximization (EM) algorithm and a k-means clustering algorithm. Two implementations of the EM algorithm (*mclust*, Scrucca et al. (2016), and *Rmixmod*, Lebret et al. (2015)) give very similar results, k-means (as implemented in *kmeans* function base *R*), however, shows only the repulsive effect. This is because k-means is a ‘hard clustering’ algorithm that cannot always assign the observations to one stimulus or the other, so ‘averaging’ solutions like the one shown in Figure 3B (middle) are not possible. This approach is similar to our toy example and is suboptimal because the uncertainty in assignments is not accounted for.

**Figure S4:**
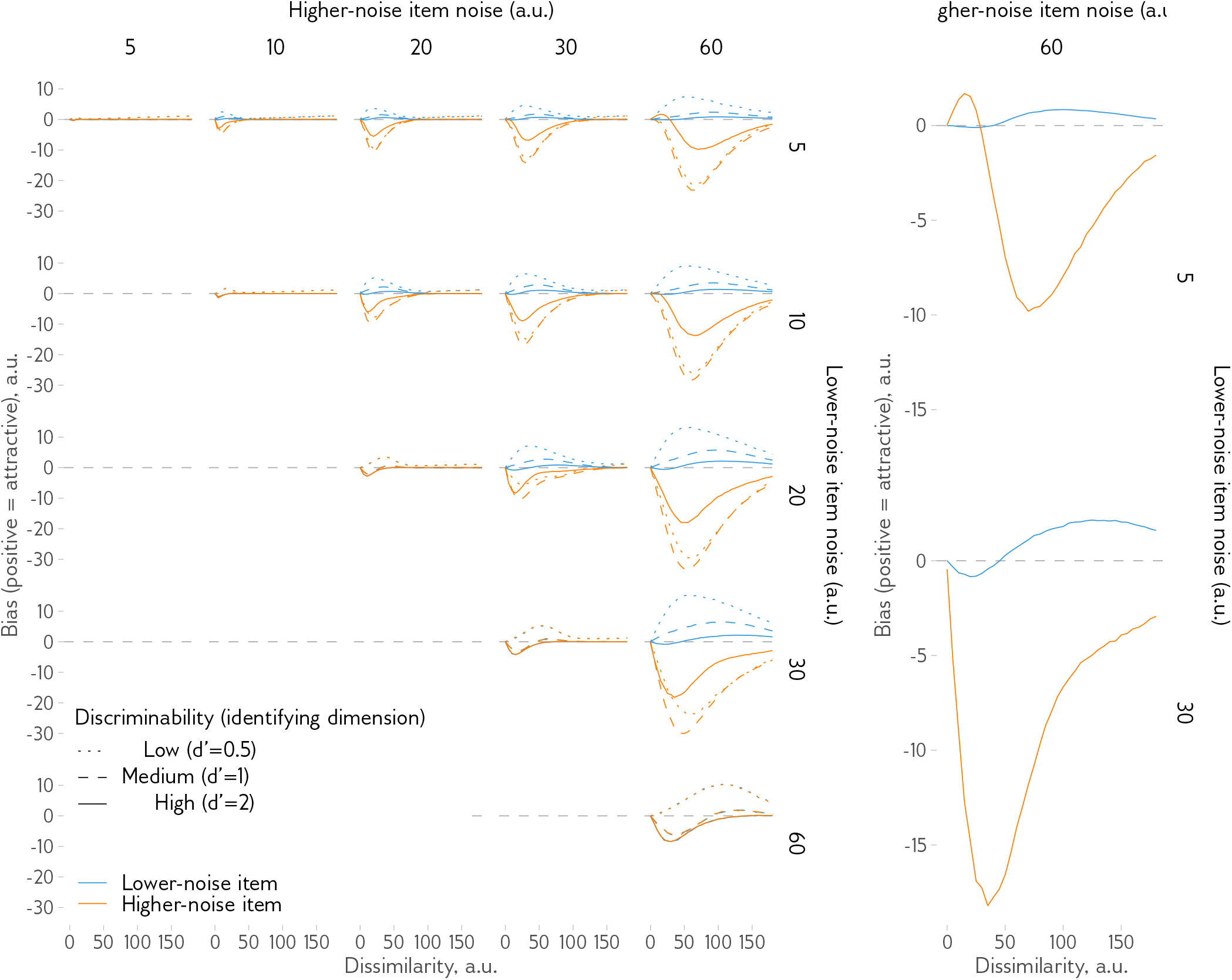
Predicted bias patterns for different feature noise levels and discriminability on the identifying dimension. Each line shows an average bias as a function of dissimilarity between stimuli (on the response dimension), with different discriminability (on the identifying dimension) indicated by the line type, and colours indicate the amount of feature noise on the response dimension for each of the stimulus. Diagonal panels correspond to Figure 4B. Medium discriminability in the off-diagonal panels correspond to Figure 5B. In general, the divergent pattern of biases shown in Figure 5B is also observed at other levels of discriminability but in some cases, highlighted for the high-discriminability condition in two plots on the right, the pattern can be different.

**Figure S5:**
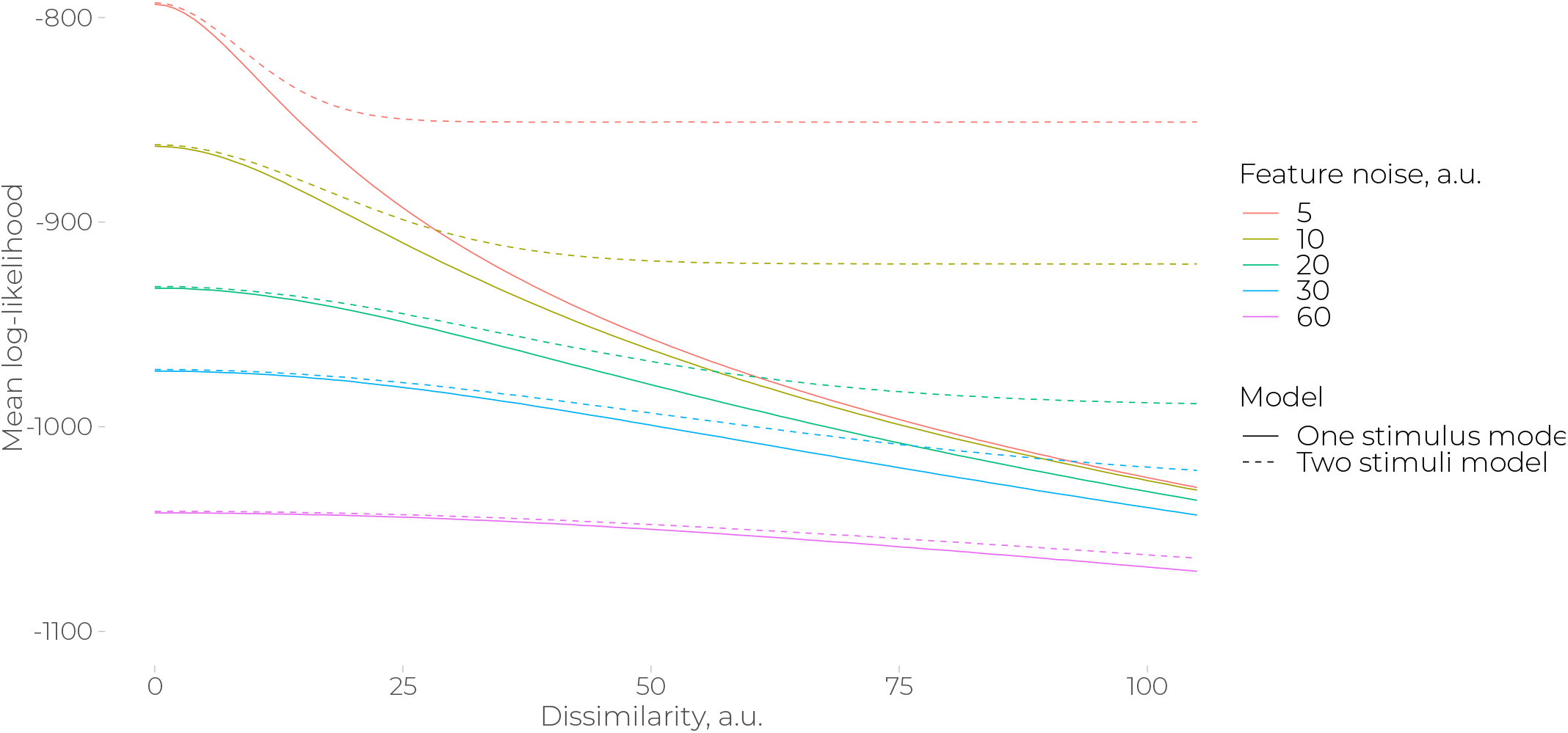
Log-likelihood of a single-stimulus and a two-stimuli models. The two-stimuli model is always more likely than a single-stimulus one since the latter is a special case of the former.

**Figure S6:**
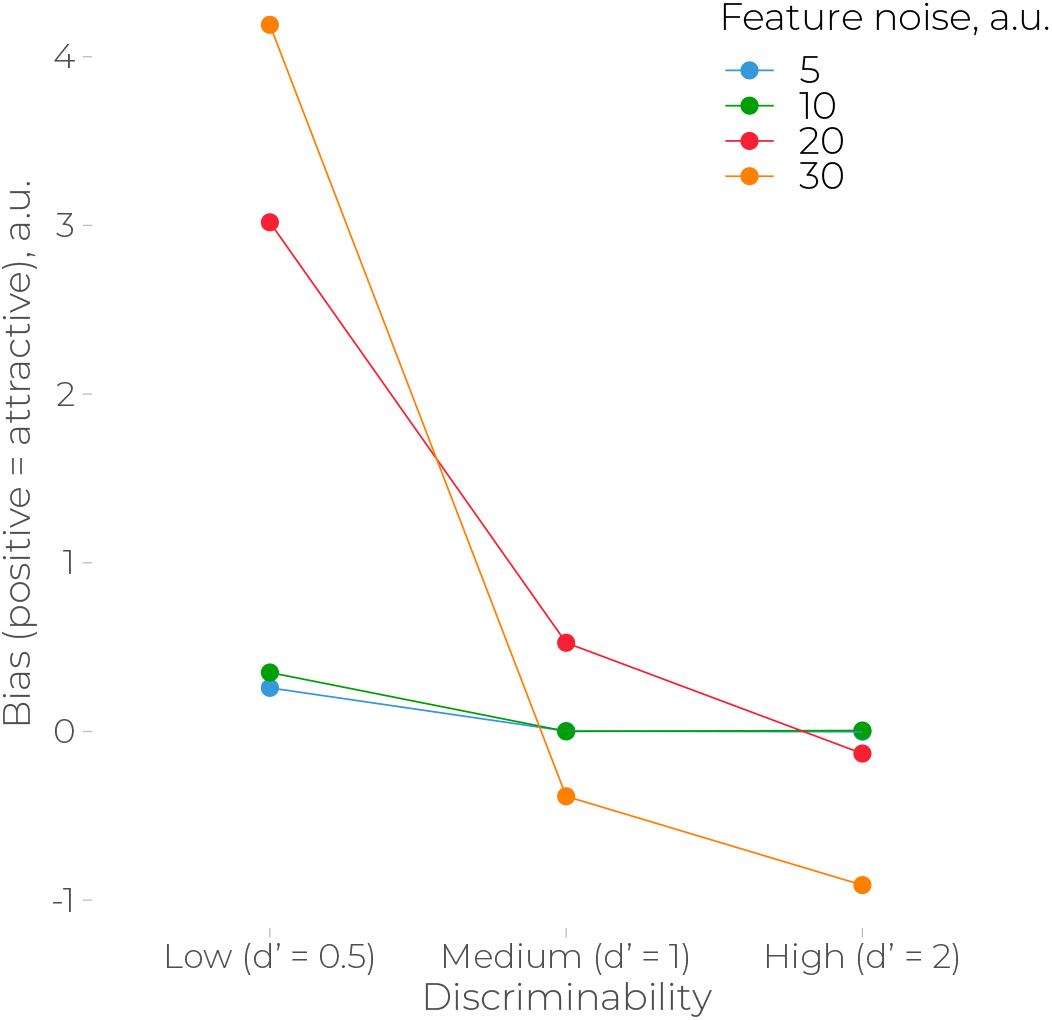
Average bias as a function of discriminability on the identifying dimension and feature noise. The data is the same as in Figure 4B but plotted only for 40 a.u. feature dissimilarity to illustrate the effect of discriminability at different feature noise levels.

## Footnotes

1 Here, I refer to serial dependence as perceptual effect for simplicity, although it also involves post-perceptual processes (see Pascucci et al. 2023 for review).

2 Bae and Luck (2017) suggested a relational coding approach for repulsive biases in working memory with simultaneously presented items. It is inspired by the idea of reference repulsion and can be seen as a version of efficient coding models. Note, however, that this is not a normative but a descriptive model, since it does not explain why such coding occurs. Furthermore, the model involves circularity in explanations (a stimulus has to be already encoded to serve as a reference, but then it should not be biased) and would be difficult to implement computationally (e.g., a single item then should create a repulsion from itself).

3 Note also how the amount of noise is underestimated in Figure 3B, left panel. This is because of the same reason: similar observations have a higher probability of coming from a given source than dissimilar ones. This means, in turn, that their estimated variability decreases.

